# Retinoic acid-dependent loss of synaptic output from bipolar cells impairs visual information processing in inherited retinal degeneration

**DOI:** 10.1101/2023.10.29.564225

**Authors:** Logan Ganzen, Shubhash Chandra Yadav, Mingxiao Wei, Hong Ma, Scott Nawy, Richard H Kramer

## Abstract

In retinitis pigmentosa (RP), rod and cone photoreceptors degenerate, depriving downstream neurons of light-sensitive input, leading to vision impairment or blindness. Although downstream neurons survive, some undergo morphological and physiological remodeling. Bipolar cells (BCs) link photoreceptors, which sense light, to retinal ganglion cells (RGCs), which send information to the brain. While photoreceptor loss disrupts input synapses to BCs, whether BC output synapses remodel has remained unknown. Here we report that synaptic output from BCs plummets in RP mouse models of both sexes owing to loss of voltage-gated Ca^2+^ channels. Remodeling reduces the reliability of synaptic output to repeated optogenetic stimuli, causing RGC firing to fail at high stimulus frequencies. Fortunately, functional remodeling of BCs can be reversed by inhibiting the retinoic acid receptor (RAR). RAR inhibitors targeted to BCs present a new therapeutic opportunity for mitigating detrimental effects of remodeling on signals initiated either by surviving photoreceptors or by vision-restoring tools.

**Significance Statement:** Photoreceptor degenerative disorders such as retinitis pigmentosa (RP) and age-related macular degeneration (AMD) lead to vision impairment or blindness. Vision mediated by surviving photoreceptors or artificial vision restoration technologies, rely on bipolar cells retaining normal function despite photoreceptor death. We find that in two animal models of RP, synaptic transmission from both rod and cone bipolar cells is severely impaired owing to diminished voltage-gated calcium current, preventing postsynaptic amacrine cells and retinal ganglion cells from properly receiving and encoding visual information. We find that an inhibitor of the retinoic acid receptor restores both the calcium current and synaptic release from bipolar cells. These discoveries about bipolar cells reveal a new functional deficit in blindness and a potential therapeutically important solution.

## Introduction

Rod and cone photoreceptors degenerate in retinitis pigmentosa (RP) and age-related macular degeneration (AMD), depriving the visual system of light-evoked signals necessary for sight. The remaining retinal neurons can survive long after the rods and cones are gone. Critically, retinal ganglion cells (RGCs), the output neurons of the retina, remain connected to the brain (Medeiros and Curcio, 2001; Mazzoni et al., 2008). This enables responses initiated by surviving photoreceptors to remain detectable late into disease progression (Ellis et al., 2023). The maintained connectivity of RGCs provides a lifeline for restoring vision in profound blindness, either by regenerating lost photoreceptors from stem cells, or by conferring artificial light sensitivity into surviving retinal neurons with optoelectronic (Humayun et al., 2012), optogenetic (Bi et al., 2006; Sahel et al., 2021), or opto-pharmacological tools (Polosukhina et al., 2012; Tochitsky et al., 2014). Whether direct or indirect, if a light stimulus can alter RGC firing appropriately, the response should be transmitted to the brain and result in visual perception.

Unfortunately, downstream neurons undergo remodeling that can corrupt the proper encoding of information and exacerbate vision loss (Telias et al., 2022). Morphological remodeling can require months to develop in mice (Pfeiffer et al., 2020), while, in contrast, physiological remodeling begins almost immediately when photoreceptors die (Stasheff et al., 2011; Telias et al., 2019). Many RGCs become hyperactive, spontaneously firing bursts of action potentials that obscure responses initiated by surviving photoreceptors. Patch clamp recordings show that the intrinsic electrical properties of RGCs are altered by photoreceptor degeneration (Tochitsky et al., 2014) and transcriptional analysis shows altered expression of voltage-gated ion channels (Tochitsky et al., 2016). In addition, the plasma membrane of RGCs exhibits increased permeability to large dye molecules and azobenzene photoswitches, events attributed to increased expression of ATP-gated P2X receptors, which have a large pore (Tochitsky et al., 2016).

How does photoreceptor death in the outer retina lead to physiological remodeling of RGCs in the inner retina? We found that retinoic acid (RA) is the key signal that triggers RGC hyperactivity and hyperpermeability (Telias et al., 2019). Inhibitors of retinaldehyde dehydrogenase (RALDH), the enzyme that synthesizes RA, or RAR, the nuclear retinoic acid receptor, reduce hyperactivity and hyperpermeability. These agents also unmask responses of RGCs to dim light, sharpen responses of visual cortical neurons to patterns of illumination, and improve behavioral visual perception in vision-impaired mice with partial photoreceptor degeneration (Telias et al., 2022).

Bipolar cells (BCs) are located between photoreceptors and RGCs. During photoreceptor degeneration, synapses onto BCs exhibit morphological remodeling. BC dendrites, which normally invaginate the synaptic terminals of rods and cones, begin to retract (Marc et al., 2003), disconnecting the input synapse. Synaptic input is also impaired by the loss or redistribution of dendritic mGluR6 receptors (Dunn, 2015) and TRPM1 channels (Gayet-Primo and Puthussery, 2015), which normally underlie the ON-BC response to the photoreceptor neurotransmitter glutamate. In addition, the expression of certain voltage-gated potassium channels is misregulated in degenerated retina (Schilardi and Kleinlogel, 2022; Schilardi et al., 2023). Despite these negative consequences, the retina shows resilience by employing mechanisms that partially compensate for photoreceptor loss. Over time, new dendrites sprout from BCs and can re-establish synaptic connections in retinas with degenerated photoreceptors (Lin et al., 2012). Experimental ablation of half of the rods in adults rebalances excitatory and inhibitory synaptic strength to BCs to restore proper RGC responses (Care et al., 2019). In young mice, partial ablation of cones leads to dendritic rewiring in certain BCs to restore the original number of input synapses (Shen et al., 2020).

In contrast, the question of whether BC output synapses are changed by photoreceptor degeneration has remained unanswered. This is critical, as changes in BC output will alter the response properties of downstream RGCs, the only cell type that provides visual input to the brain. Here we evaluate the effects of photoreceptor degeneration on output synapses of two types of ON-BCs; rod bipolar cells (RBC), critical for scotopic vision and the most numerous BC type in mouse, and cone bipolar cell type-6 (CBC6), which plays a crucial role in all vision, from scotopic, to mesopic, to photopic. We find a dramatic reduction of voltage-gated Ca^2+^ current and neurotransmitter release from both RBCs and CBC6s, weakening synaptic output and decreasing the frequency response of the BC to RGC synapse. These effects can be reduced by inhibiting RA signaling, providing a potential therapeutic approach for abrogating the detrimental effects of BC remodeling in RP and AMD.

## Materials and Methods

### Animals

Mice were handled in accordance with protocols approved by the UC Berkeley Institutional Animal Care and Use Committee (AUP-2016-04-8700-1) and conformed to the NIH Guide for the Care and Use of Laboratory Animals. C3H/HeJ mice (Jackson 000659) were used as the rd1 model (*Pde6b^rd1^*). B6.CXB1-Pde6b^rd10^/J mice were used as the rd10 model (Jackson 004297). CCK-ires-Cre knock-in mice (Jackson 012706, abbreviated CCK-Cre) were used to drive Cre recombinase under the cholecystokinin promoter to label CBC6 cells. BAC-Pcp2-IRES-Cre mice (Jackson 010536, abbreviated Pcp2-Cre) were used to drive Cre recombinase under the Pcp2 promoter to label RBCs. Ai32 mice (Jackson 024109, B6.Cg*-Gt(ROSA)26Sor^tm32(CAG-COP4*H134R/EYFP)Hze^*/J) were used to drive channelrhodopsin-2/eYFP fusion protein in Cre-expressing CBC6 or RBC cells. Both rd1 and rd10 mice strains were homozygous for their respective mutations in the *Pde6b* locus. The CCK-Cre, Pcp2-Cre, and Ai32 alleles were always used in mice in a hemizygous state. Mice of both sexes were used interchangeably. For experiments with WT and rd1 mice, animals were used within 3 days of postnatal day 60 (p60). For experiments with rd10 mice, animals were used within 3 days of p90.

### Retinal Dissection

Mice were euthanized via isoflurane exposure and internal decapitation. Retinal dissection was performed with enucleated eyes in oxygenated ACSF (in mM: 119 NaCl, 26.2 NaHCO3, 11 Dextrose, 2.5 KCl mM, 1 K2HPO4, 1.3 MgCl2* 6H2O, 2.5 CaCl2, 293 mOsm/Kg), (5% CO_2_/95% O_2_) in room light conditions with a dissection microscope. For retinal flat-mount experiments, retinas were flattened with 4 relieving cuts. For retinal slice experiments, flattened retinas were mounted on 13 mm diameter 0.45 µm filter paper discs (MF-Millipore) and sliced to a thickness of 250 µm with a Stoelting Tissue Slicer. Retinal slices were rotated 90 degrees and mounted within a vacuum grease well.

### Electrophysiology

All RGC recordings were performed in flat-mount retinas. Recordings taken from BCs and AII amacrine cells were performed in retinal slices. For RGC recordings, retinas were isolated and incubated in ACSF containing hyaluronidase (9,800 U/mL) and collagenase (2,500 U/mL) for 10 minutes to facilitate penetrating of the inner limiting membrane allowing access to RGC somas for patch clamp recording (Schmidt and Kofuji, 2011). Retinas were mounted ganglion cell side up in a recording chamber and held in place with a harp (Warner Instruments), superfused with ACSF oxygenated with 5% CO_2_/95% O_2_ at a rate of 5 ml/minute at 34 degrees C and viewed under DIC optics with an upright microscope (Olympus). Reagents for ACSF were purchased from Fisher Scientific. ON α-RGCs did not visually express ChR2-eYFP in the CCK-ires-Cre mouse x Ai32 mouse lines. EPSCs from ON α-RGCs were completely blocked with the AMPAR antagonist DNQX. EPSCs also had a delay from light flash to onset of about 6 ms. Other RGCs, that were not ON α-RGCs, expressed ChR2-eYFP that exhibited near instantaneous current with light flash with a delay of about 0.2 ms which was insensitive to DNQX.

For retinal slice experiments, slices were made as described and perfused in the same way as flat-mount retinas. For direct recording from RBCs, ACSF CaCl2 was substituted with equimolar BaCl_2_ to prevent Ca^2+^-activated chloride currents from ANO1 channels (Paik et al., 2020). Electrodes were made with boroscilicate capillary glass tubing with OD = 1.5 mm, ID = 1.17 mm (Warner G150TF-4). For RGC recording, glass was pulled to a resistance of 5-6 MΩs using a Narishige pipette puller. For BCs and amacrine cell recording, electrodes were made with a resistance of 7-8 MΩs with a Sutter Instruments P1000. For voltage clamp experiments, the pipette solution contained (in mM): 123 Cs gluconate, 8 NaCl, 1 CaCl_2_, 10 EGTA, 10 HEPES, 10 glucose, 5 Mg^2+^ ATP, 5 QX 314 (Tocris Bioscience), 0.01 Alexa 594, pH 7.4 with CsOH, 293 mOsm/Kg. For current clamp experiments in BCs to measure optogenetic responses, the pipette solution contained (in mM): 125 K^+^ Gluconate, 10 EGTA, 10 KCl, 10 HEPES, 4 Mg^2+^ ATP, 0.01 Alexa 594, 293 mOsm/Kg. Alexa Fluor 594 Hydrazide (ThermoFisher Scientific) at 10 µM was used to dye fill all cells during recording for confirmation of cell identity. Whole-cell recordings were sampled at 20 kHz and filtered at 2 kHz with a Multiclamp 700B amplifier, digitized with a 1440a or 1550a Digidata A-D converter and analyzed offline with Axograph X or Clampfit. Recordings with series resistance above 10% of the cell membrane resistance or reached above 30 mΩ were discarded.

To block photoreceptor responses to light, the kainate receptor antagonist ACET (1 µM) and the mGluR6 agonist L-AP4 (10 µM) were added to ACSF. Gabazine (10 µM) and strychnine (10 µM) were also added to block ionotropic GABA and glycine receptors. MFA (100 µM) was used to inhibit gap junctions. Pharmacological agents were purchased from Tocris Biosciences.

Optogenetic stimulation of CBC6s and RBCs was provided by a 470 nm LED delivering 2.1X10^8^ photons/s, (Lumencore or CoolLED pE-4000) measured at the plane of the retina with a spectrophotometer (Thor Labs). For Fig. 2 and 6, stimulation intensity was varied by changing the duration of the stimulus from 0.1 to 10 ms. Thus, the number of photons delivered during the optogenetic stimulus ranged from 2.1X10^7^ (100%) to 2.1X10^5^ (1%).

**Figure 1.**
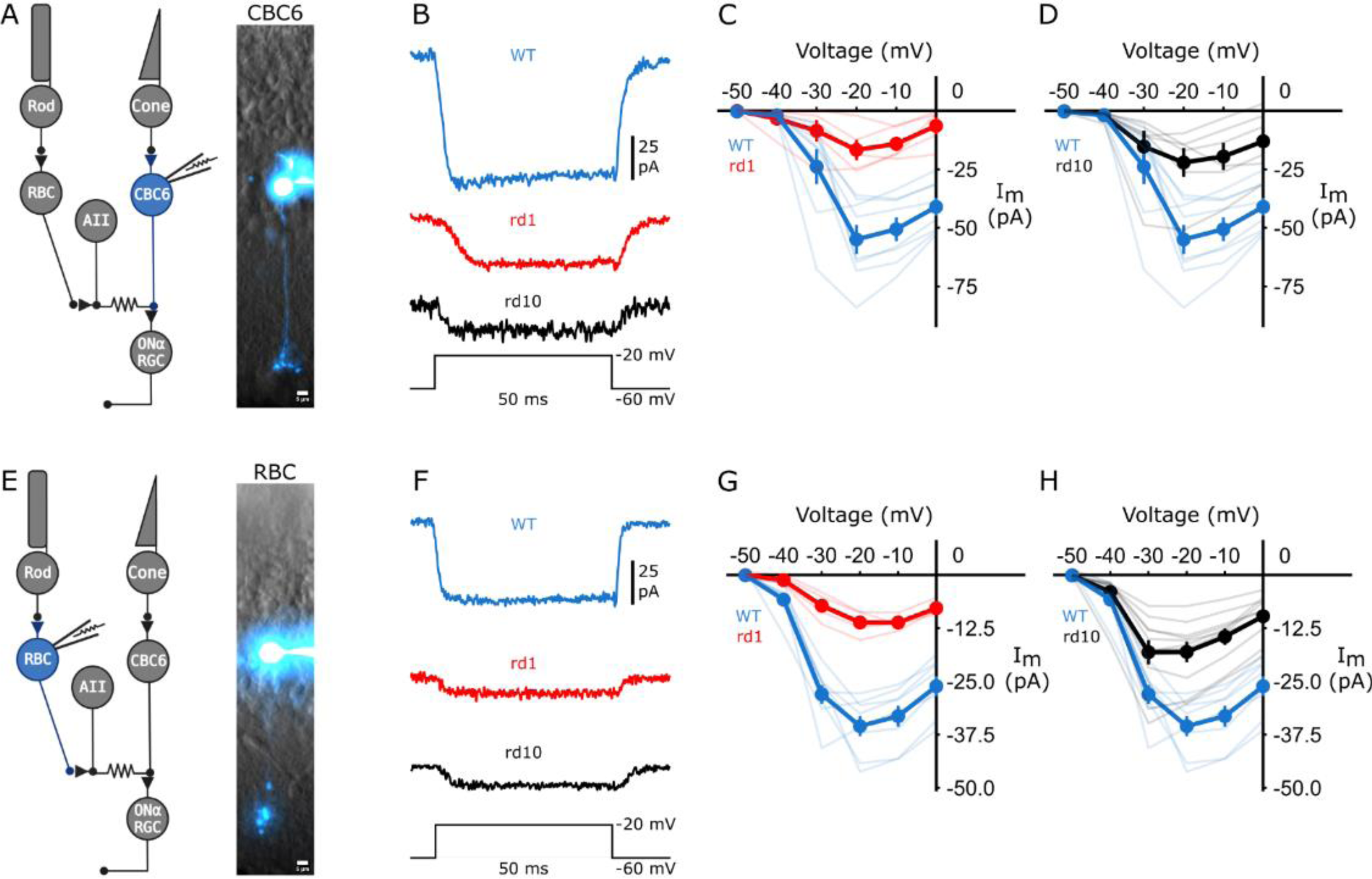
Voltage-gated calcium current of ON-BCs is reduced in photoreceptor degenerated retinas. A, E) Retinal circuit diagrams showing whole-cell patch-clamped CBC6 cells or RBCs with accompanying fluorescent dye fills (Alexa-Fluor 594) of a CBC6 cell or an RBC in a retinal slice. Scale bars = 5 µm. B, F) Inward calcium currents in a CBC6 and RBC from WT, rd1, and rd10 retinas. Current was activated with a depolarizing voltage step (50 ms) from −60 mV to −20 mV. C, D) Steady-state current vs. voltage (I-V) curves, elicited with a series of 10 mV incrementing depolarizing steps from - 60 to 0 mV from CBC6 cells WT: N = 8 rd1: N = 6; rd10: N = 6. G, H) Steady-state current vs. voltage (I-V) curves for RBCs, WT: N = 9; rd1: N = 6; rd10: N = 10). I-V curves from rd1 and rd10 BCs shows that the voltage-gated calcium current is lower than in WT BCs (Pairwise Wilcoxon Rank Sum Exact Test (PWT) with false discovery rate (FDR) correction (for CBC6: WT − rd1, p = 0.016; WT − rd10, p = 0.048. For RBC: WT − rd1, p= 0.001; WT − rd10, p-value = 0.001). Transparent traces indicate individual cells. Error bars indicate ± SEM.

**Figure 2.**
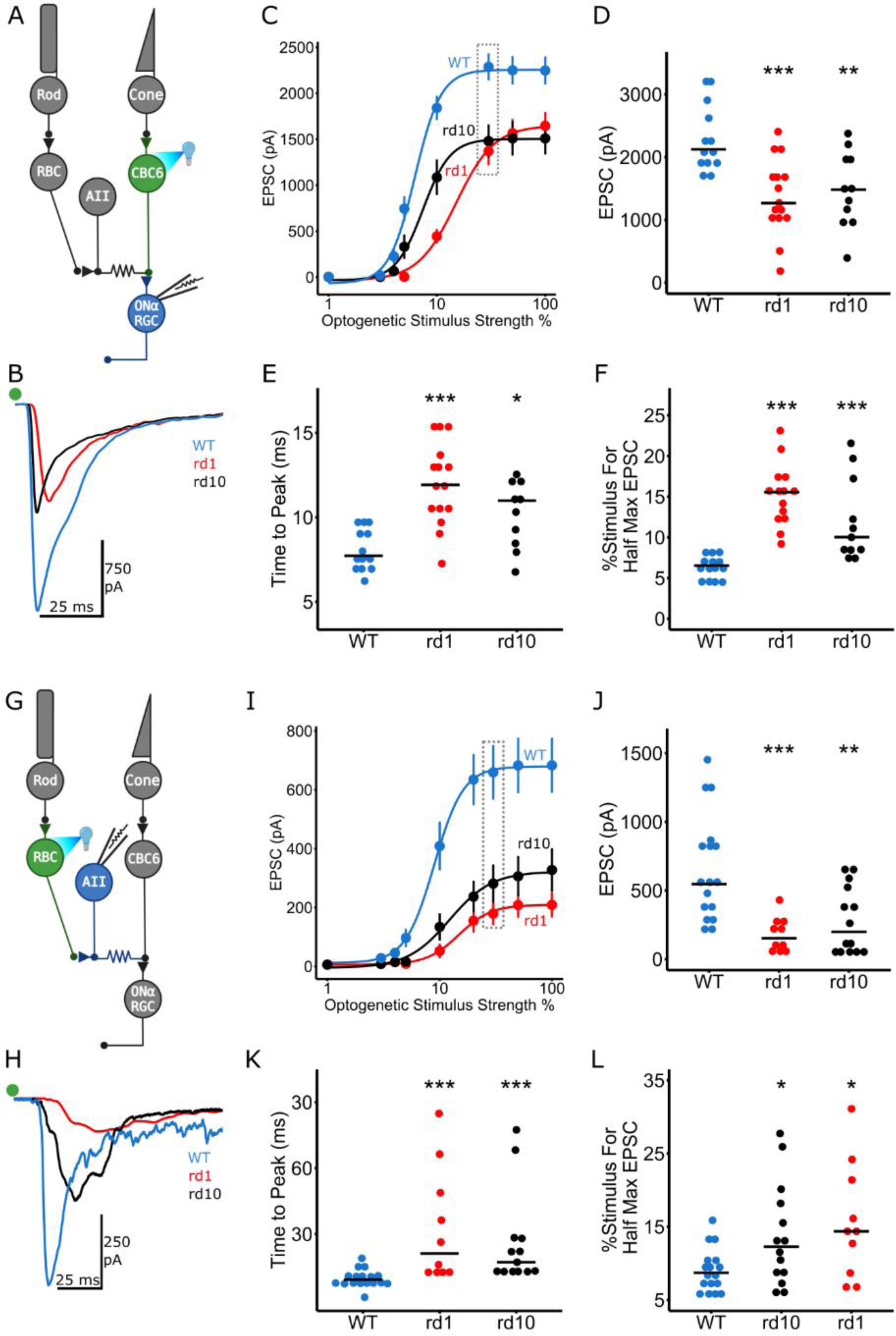
Synaptic output from ON-BCs is reduced in photoreceptor degenerated retinas. A, G) Retinal circuit diagrams showing optogenetically stimulated CBC6 cells or RBCs. EPSCs were recorded from a postsynaptic ON α-RGC or AII amacrine cell, respectively. B,H) EPSCs in CBC6 cells (B) and RBCs (H). Note the reduced EPSCs in rd1 and rd10 retinas compared to WT. The green dot represents the light flash. C,I) At all effective optogenetic stimulus strengths, EPSCs from CBC6 cells (C) and RBCs (I) were smaller in rd1 and rd10 than in WT (For CBC6 cells: rd1 vs. WT: p<0.001; rd10 vs. WT: p = 0.009; rd1:N=15, rd10:N=11, WT:N=13. For RBCs: rd1 vs. WT: p<0.001; rd10 vs. WT: p<0.001; WT:N=17, rd1:N=10, rd10:N=14). D,J) EPSCs elicited by a saturating stimulus (dashed box in panels C and I) were smaller in rd1 and rd10 than in WT (For CBC6 cells: rd1= 1.27 nA, rd10 = 1.48 nA, WT = 2.12 nA; rd1 vs. WT: p <0.001; rd10 vs. WT: p = 0.008; for RBCs: rd1 = 152 pA, rd10 = 198 pA, WT = 546 pA; rd1 vs. WT: p <0.001; rd10 vs. WT: p = 0.007). E,K) Rise time of EPSCs is slower in rd1 and rd10 than in WT retinas for both the CBC6 output synapse (rd1 = 11.92 ms, rd10 = 10.99 ms, WT = 7.72 ms; rd1 vs. WT, p <0.001; rd10 vs. WT, p = 0.011) and the RBC output synapse (rd1 = 21.17 ms, rd10 = 19.19 ms, WT = 9.24 ms; rd1 vs. WT, p <0.001; rd10 vs. WT, p <0.001). F,L) The calculated stimulus strength required to elicit half-maximal EPSCs from CBC6 cells (F) and RBCs (L) is higher for rd1 and rd10 than for WT. (For CBC6 cells: rd1= 15.5%, rd10 = 10.0%, WT = 6.5%; rd1 vs. WT: p <0.001; rd10 vs. WT: p <0.001; for RBCs: rd1 = 14.3%, rd10 =12.3%, WT = 8.7%; rd1 vs. WT: p = 0.02; rd10 vs. WT: p = 0.05). For panels D – F, and J – L, each data point represents EPSC statistics from an individual recording; horizontal lines represent median EPSC values. Error bars indicate ± SEM. All statistical comparisons were made with PWT with FDR correction.

### Mapping ON α-RGC Virtual Spatial Receptive Rield

Spots of light with a 50 µm diameter were generated with a Texas Instruments Digital Light Projector (CEL1015 Light Engine, Digital Light innovations) controlled with CELconductor Control Software. Briefly, an ON α-RGC was identified and patch clamped as described above. The cell body was focused on with an Olympus LUMPlanFl 20x/0.5 W ∞/0 objective and centered within a 650 µm x 650 µm field of view. Each spot projection and simultaneous EPSC recording was controlled via Digidata.

### Trans-Retinal Stimulation

To achieve electrical trans-retinal stimulation, a custom chamber was created. A 65 mm petri dish was shaved down in height and embedded within a 100 mm petri dish with RTV sealant. The 65 mm dish was elevated ∼ 3 mm from the bottom of the 100 mm dish. An approximately 1 mm hole was melted in the center of the 65 mm dish with a 21 gauge needle. To mount a retina, a ∼ 7.5 mm x 7.5 mm piece of ion-permeant dialysis membrane was placed over the 1 mm hole to supply an even surface to place a flat-mount retina, RGC side up. A ∼ 7.5 mm x 7.5 mm piece of flexible cellophane with an ∼ 1mm hole punched into the center was placed over the retina as a coverslip. The retina is then sealed between the membrane and coverslip with vacuum grease so that the only electrical path between the two dishes is through the retina. Two copper wires were placed in each dish below and above the retina which were connected to an ISO-Flex stimulus isolation unit. Electrical stimulation was achieved by setting the desired voltage on the ISO-Flex and triggering the stimulus via Digidata.

### 2-Photon iGluSnFR Imaging

2-Photon fluorescence imaging was performed on a Sutter Instruments moveable objective microscope with an Olympus XLUMPlanFl 20x/0.95 W ∞/0 objective. iGluSnFR excitation was achieved with a Coherent Chameleon Ultra II tuned to 910 nm. Imaging was performed with ScanImage software. Scanning field of view was 25 µm x 25 µm. Images collected in Fig 4c-d were scanned at 256 pixels per line x 256 lines (frame rate = 1.48 Hz, 2 ms per line). Higher time resolution of iGluSnFR flouresence for Fig 4e-g was performed at 256 pixels per line x 6 lines (frame rate = 63.13 Hz, 2 ms per line).

### Immunohistochemistry

ON α-RGCs were injected with Alexa Fluor 568 hydrazide (A10441, Thermo Fisher) as described previously (Tetenborg et al., 2017). In short, somas of RGCs in the flat-mount retina were visualized by acridine orange labeling (Meyer et al., 2014). Potential ON α-RGCs were identified based on their large soma size (>20 µm) and targeted with sharp microelectrodes containing Alexa Fluor 568. The dye injected retinas were fixed with 4% PFA and the following immunolabeling was carried out as described before (Yadav et al., 2019). In brief, the flat-mount retinas were incubated into primary and secondary antibodies for 3 and 1 days, respectively. CtBP2 (mouse, AB_399431, BD Biosciences) monoclonal and secretagogin (sheep, AB_2034062, BioVendor) polyclonal antibodies were used at a dilution of 1:500 and 1:750, respectively. All immunostainings were performed at room temperature. Images were scanned with 40x/1.4 oil objective using Zeiss LSM780 Laser Scanning Confocal Microscope. The z-stacks were acquired at a constant pixel size of 60nm*60nm and a z-step of 0.3µm unless stated otherwise. All raw images thus obtained were deconvolved with Huygens Essential software (Scientific Volume Imaging, Netherlands). The deconvolution was achieved by using a theoretical point spread function. The resulting images were further processed and analyzed in Fiji (https://fiji.sc/) (Schindelin et al., 2012). The z-stacks were normalized using stack histograms prior to analysis. The colocalization analysis was carried out within each optical section unless stated otherwise. Brightness and contrast of the images were adjusted post-analysis in Fiji.

Colocalization between channels was performed with Fiji utilizing the colocalization highlighter plugin as previously described (Behrens et al., 2022). Briefly, deconvoluted z-stacks were segmented using stack histogram. Secretagogin and CtBP2 signals were thresholded with Otsu’s method, and dye injected RGCs were manually thresholded. With colocalization highlighter, an intensity ratio of 50% was applied to estimate colocalization between Secretagogin and CtBP2. Colocalization of Secretagogin and CtBP2 with RGCs was performed sequentially with an intensity ratio of 10%.

Resultant 8-bit images of puncta were generated where at least 5 or more connected voxels were considered as colocalization. These images were overlayed with either RGCs or CBC6 axon terminals, and the 3D Objects Counter plugin was used to quantify putative synaptic contacts. To isolate individual CBC6 axon terminals, axonal endings were traced back from a common point in the z-stack 8 µm into the IPL.

### AAV Delivery of iGluSnFR

Plasmid containing iGluSnFR(A184S) under the control of the human synapsin promoter was purchased from Addgene (#106174) and packaged in the AAV8-Y733F serotype (Dalkara et al., 2013). Viral constructs were generated in the Gene Delivery module of the Vision Science Core at UC Berkeley. One month prior to imaging, 1.5 μl of virus (1×10^15^ vg/mL) was injected into intravitreally into the eye of rd1 or WT mice.

### Data Visualization and Statistics

All reported statistics and diagrams were generated with R 4.2.2. Retinal and experimental diagrams were created with Biorender.com.

## Results

### Voltage-gated calcium current in ON-BCs is reduced by photoreceptor degeneration

Calcium influx through voltage-gated Ca^2+^ channels is critical for evoking neurotransmitter release at the BC output synapse (Tachibana et al., 1993). To measure the voltage-gated Ca^2+^ current, we recorded from BC somata in retinal slices, a preparation that made the cells easy to identify and access with a patch pipette. After establishing whole-cell voltage clamp we applied a series of depolarizing voltage steps. We compared activation of the voltage-gated Ca^2+^ current in BCs from wildtype (WT) mice from those in rd1 and rd10 mice, widely used animal models of RP. We focused on the Type-6 ON cone bipolar cell (CBC6), which synapses directly onto RGCs, and rod bipolar cells (RBCs), which drive RGCs indirectly through a serial synapse onto the CBC output synaptic terminal (Fig. 1A,E). Candidate BCs of each type were identified with cell-type selective gene promoters that drive expression of a fluorescent protein reporter fused with channelrhodopsin-2 (ChR2-eYFP), namely the Cck promoter for CBC6 cells and the Pcp2 promoter for RBCs. The identity of individual BCs was confirmed by dye-filling the cell to localize their terminals to specific sublamina in the inner plexiform layer (IPL) (Fig. 1A,E).

We found that the voltage-gated Ca^2+^ current in CBC6 cells was 3-fold larger in WT mice than in rd1 or rd10 mice at maximal activation (Fig.1B–D). Similarly, voltage-gated Ca^2+^ current in RBCs was 2-fold larger than in rd1 or rd10 mice at maximal activation (Fig. 1F–H). In both BC types, steady-state current vs. voltage curves showed reduced current amplitude without any change in voltage-dependent activation (Fig. 1C–D,G–H), consistent with a change in Ca^2+^ channel number rather than a change in gating properties or channel type. We found that the L-type Ca^2+^ channel inhibitor nimodipine eliminated the current in CBC6 cells consistent with previous findings that L-type Ca^2+^ channels predominate in BCs (Heidelberger and Matthews, 1992). Photoreceptor degeneration occurs early in postnatal development in rd1 mice and later in adulthood in rd10 mice, making certain aspects of morphological remodeling more severe in the rd1 strain (Marc et al., 2003). Despite this, the loss of Ca^2+^ current in CBC6 cells and RBCs was nearly identical in rd1 and rd10 mice.

### Transmission at the BC output synapse is reduced by photoreceptor degeneration

Neurotransmitter release is triggered by intracellular Ca^2+^, hence the reduced Ca^2+^ current in BCs suggests that synaptic transmission will also be reduced. ON-BCs are ordinarily depolarized in the light, but in the absence of photoreceptors we can use optogenetics as an alternative to depolarize the cells and evoke synaptic release. We leveraged the cell-type selective expression of ChR2-eYFP to optogenetically depolarize either RBCs or CBC6s, in the WT, rd1, or rd10 background. To interrogate the BC output synapse in isolation, we blocked photoreceptor inputs onto BCs with the mGluR6 agonist L-AP4 and the kainate receptor antagonist ACET, which together eliminate light responses in all BCs (Tien et al., 2017). We also blocked inhibitory synaptic inputs onto BCs with antagonists of ionotropic GABA and glycine receptors and electrical synapses with the gap junction uncoupler MFA to block inner retina oscillations.

We stimulated CBC6 cells with light flashes while we recorded evoked excitatory postsynaptic currents (EPSCs) from ON α-RGCs in flat-mount retinas (Fig. 2A). Amongst the various BCs, the promoter for cholecystokinin (Cck) enables transgene expression specifically in CBC6 cells, although sparse expression has been noted in non-BC cell types, including some ON type RGCs (Zhu et al., 2014). ON α-RGCs were identified by soma size and dendritic layer termination in the IPL, mapped after dye-filling. If a ChR2-eYFP expressing RGC was inadvertently targeted, it was removed from further consideration based on the exceedingly short optogenetic response delay and lack of susceptibility to the AMPAR antagonist DNQX (See Methods).

Optogenetically evoked EPSCs in ON α-RGCs were smaller and slower in rd1 and rd10 than in WT retinas (Fig. 2B). EPSCs were reduced across all stimulation strengths, including stimuli that saturated the synaptic response, consistent with diminished synaptic efficacy. EPSCs were ∼40% smaller in rd1 and rd10 retinas with a saturating stimulus (Fig. 2C,D). In addition, EPSC kinetics were slower in both rd1 and rd10, with a longer time-to-peak (Fig. 2E). The output synapse also exhibited a striking loss of sensitivity in degenerated retina. The stimulus required to evoke a half-maximal EPSC was nearly three times stronger in rd1 and rd10 compared to WT retina (Fig. 2F). Thus, the CBC6 output synapse exhibited a decrease in both peak response and sensitivity.

For examining the output synapse of RBCs, we expressed ChR2-eYFP under control of the RBC-selective promoter for Purkinje cell Protein-2 (Pcp2) (Liang et al., 2021). We optogenetically depolarized RBCs in retinal slices, recording EPSCs in their postsynaptic partner the AII amacrine cell. AII amacrine cells are electrically coupled to CBC6, indirectly driving RGC firing (Fig. 2G). We recorded EPSCs in AII cells, which were identified after dye-filling by their distinctive cell body shape and dendritic terminations in the IPL. Synaptic output of RBCs onto AII cells was also reduced in rd1 and rd10 retinas as compared to WT (Fig. 2H). Responses to saturating light at the RBC synapse were reduced by ∼75% in rd1 retinas and ∼65% in rd10 retinas as compared to WT with a saturating stimulus (Fig. 2I,J). The output synapse of RBCs also exhibited a loss of sensitivity to optogenetic stimulation, although less striking than at the CBC6 output synapse (Fig. 2L). RBCs ordinarily exhibit highly synchronous release, resulting in rapid and transient postsynaptic responses (Singer et al., 2004), but the responses were slower to rise and more prolonged in degenerated retina, implying that degeneration promotes more asynchronous transmitter release from RBCs. The time-to-peak of evoked EPSCs was 2-or 3-fold longer, for rd10 and rd1 retinas as compared to WT (Fig. 2K). In summary, the output synapses of both CBC6 cells and RBCs show dramatically reduced synaptic efficacy in degenerated retina, consistent with reduced Ca^2+^-dependent neurotransmitter release.

We were concerned that the expression of ChR2 might be lower in BCs from degenerated retinas than in BCs from WT retinas, such that the same light flash might elicit less depolarizing current resulting in a deficit in synaptic release. To test for this possibility, we used patch clamp recordings to compare the light-elicited ChR2 current in rd1, rd10, and WT retinas (Fig. 3A). The dendritic tree of BCs regresses in degenerated retinas, so to correct for the smaller surface area we measured whole-cell capacitance and calculated the current density for each cell. We found that the ChR2 current density in CBC6 cells was no different in rd1 than in WT retinas (Fig. 3B,C). Similarly, the ChR2 current density in RBCs was no different in rd10 retina than in WT (Fig. 3F,G). The kinetics of the light-elicited current were identical and there was no difference in paired currents elicited by two light flashes, suggesting no difference in activation or inactivation properties (Fig. 3 D,E). Light-elicited membrane depolarization caused by a single flash was also similar in degenerated retinas as in WT (Fig. 3H-J). It should be noted that membrane potential measurements taken from the cell body inaccurately reflect the membrane potential at the synaptic terminal, which is electrotonically distant. Because the WT cells have a larger voltage-gated Ca^2+^ current that can amplify depolarizations, the local depolarization in the BC terminals might well be larger in WT than in rd1 or rd10 terminals, even if they express the same number of ChR2 channels.

**Figure 3.**
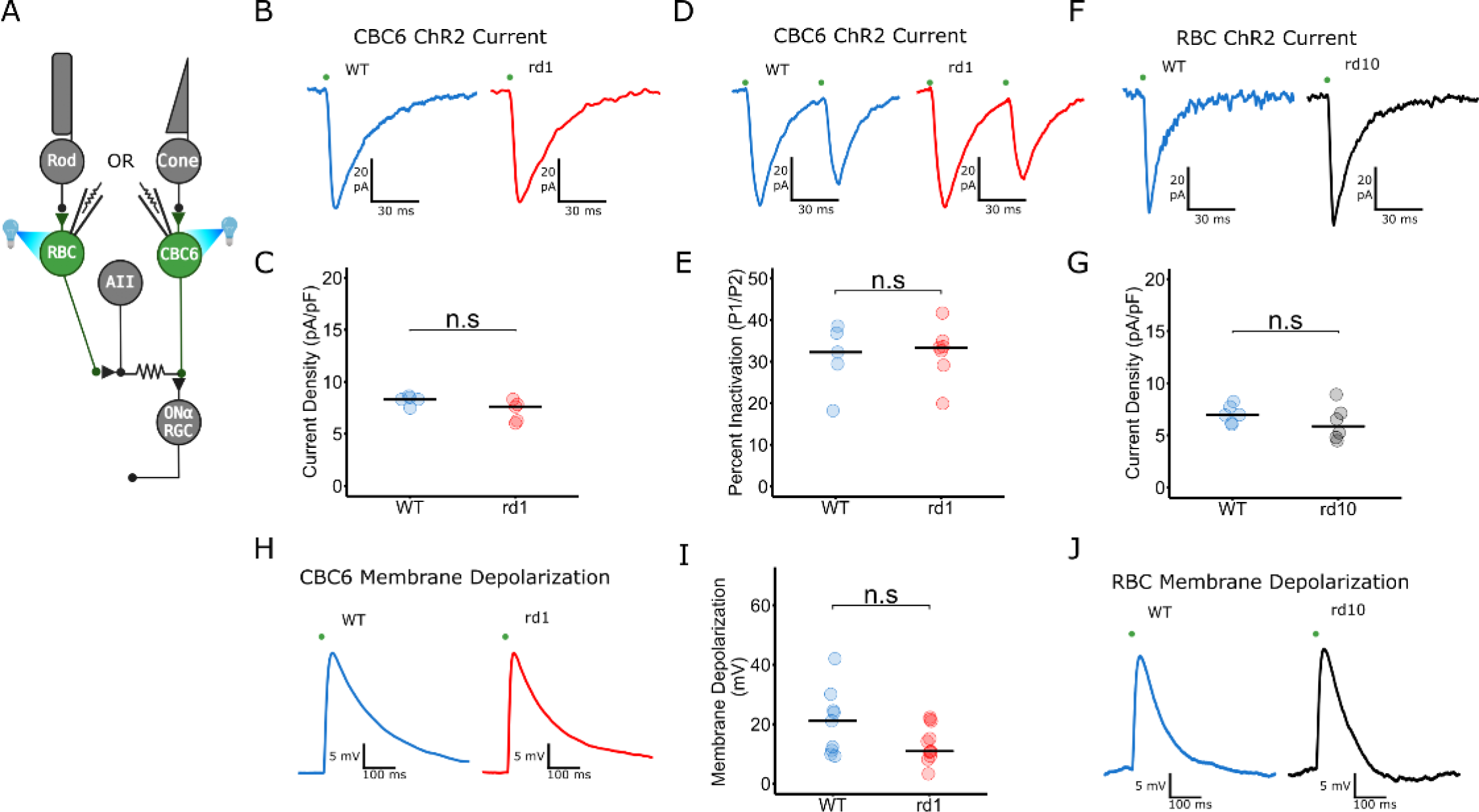
Intrinsic ChR2 function is not altered by photoreceptor degeneration. A) Retinal circuit diagram showing whole-cell patch-clamp and simultaneous optogenetic stimulation of a CBC6 cell or an RBC. B) ChR2-induced inward current elicited by optogenetic stimulation of CBC6 cells in WT and rd1. C) There is no difference in ChR2 current density in CBC6s in WT and rd1 retinas. WT: N=5, rd1: N=5 (p = 0.15). D) ChR2-induced inward current elicited in CBC6 cells by a pair of optogenetic stimuli 50 ms apart in WT and rd1. E) ChR2 current inactivated approximately 30% in the second flash with no difference between both rd1 and WT retinas. WT: N=5, rd1: N=7 (p = 0.88). F) ChR2-induced inward current elicited by optogenetic stimulation of RBCs in WT and rd10. G) There is no difference in ChR2 current density in RBCs in WT and rd10 retinas WT: N=6, rd1: N=6 (p = 0.35). H) ChR2-induced membrane depolarization in a WT and rd1 CBC6 cell. I) There was no difference in the ChR2-elicited membrane depolarization. WT: N=9, rd1: N=11 (p = 0.11). J) ChR2-induced membrane depolarization in a WT and rd10 RBC. CBC6 optogenetic pulse width: 2.5 ms. RBC optogenetic pulse width: 1 ms. Horizontal lines represent median values. All statistical comparisons were made with PWT with FDR correction.

### Morphological analysis shows that the number of output synapses from CBC6 cells is unaltered by photoreceptor degeneration

Reduced synaptic transmission could either be caused by a decrease in the number of synaptic contacts between pre- and postsynaptic neurons or by a decrease in the synaptic efficacy with no change in synapse number. To distinguish between these possibilities, we used immunohistochemistry and fluorescence microscopy to quantify the density of CBC6 synaptic contacts onto ON α-RGC dendrites in the IPL. We used an antibodiy against secretagogin to visualize the axon terminals of CBC6 cells. Secretagogin labels CBC types 2,3,4, 5, 6 in the mouse retina. CBC2-4 are OFF-BCs that terminate in distant OFF sublamina, while CBC5 stratifies in IPL 3, so CBC6 terminals can be identified unambiguously in IPL 4 (Puthussery et al., 2010). CBC6 cells constitute more than 70% of bipolar synaptic inputs to ON α-RGC (Schwartz et al., 2012) while other ON-CBC types sharing stratification with ON α-RGCs lack secretagogin expression (Helmstaedter et al., 2013). We labeled BC synaptic ribbons with a fluorescent antibody against ribeye, the main structural protein in the ribbon. Finally, we visualized the dendritic tree of an individual ON α-RGC by dye-filling the cell (Fig. 4A).

**Figure 4.**
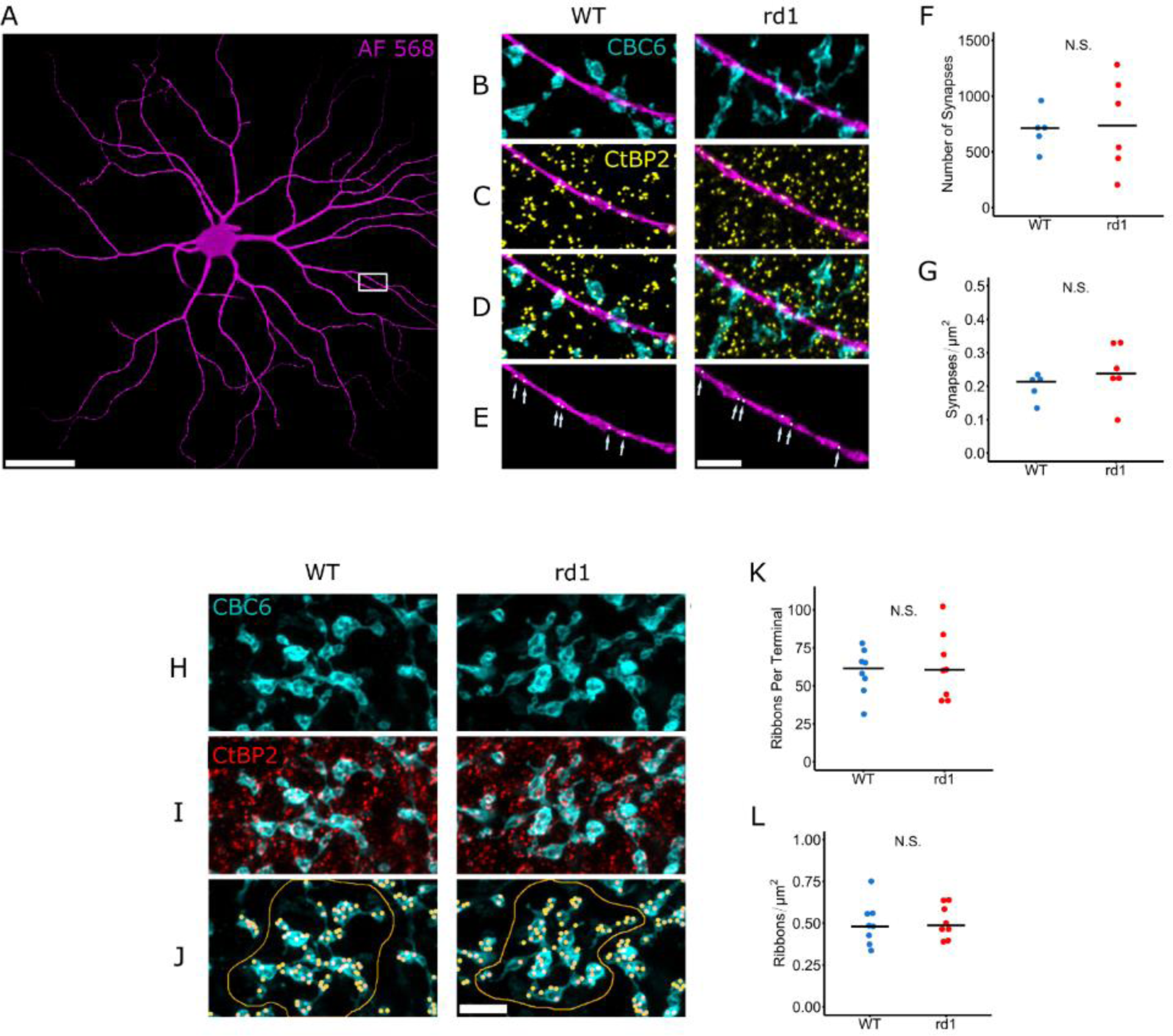
CBC6 cell output synapse morphology is unaltered by photoreceptor degeneration. A) Soma and dendrites of an ON α-RGC, labeled by intracellular injection with Alexa Fluor 568. Image is a 3-D projection from a stack of confocal optical sections. B-E) Magnified view of one ON α-RGC dendrite (magenta), overlapping with CBC6 axon terminals (B), labeled with anti-secretagogin (cyan), C) overlapping with synaptic ribbons, labeled with anti-CtBP2 (yellow), D) overlapping with synaptic ribbons exclusively in CBC6 terminals, identified by co-labeling with anti-secretagogin and anti-CtBP2. E) Arrows point to sites of co-localization between the dendrite, CBC6 axon terminals, and CtBP2, i.e., presumptive ribbon synapses between CBC6 cells and the ON α-RGC. F, G) Quantification of the total number of CBC6 ribbon synapses across all optical sections of the dendritic tree of an ON α-RGC (F) and the density of CBC6 ribbon synapses per unit area of the ON α-RGC. (G). There is no significant difference in the number (WT vs. rd1: p = 1) or density of ribbon synapses (WT vs. rd1: p = 0.33) in rd1 vs. WT retinas. WT: N=5, rd1: N=6. H)2D projection of the 3D structure of axon terminals of CBC6 cells in WT and rd1 retinas. I) CBC6 axon terminal overlapping with CtBP2. J) CBC6 axon terminals with yellow dots denoting thresholded points of overlap with CtBP2. The red outline indicates the axon terminal tree of one individual CBC6 cell. K, L) Quantification of the number of CtBP2 sites per CBC6 axon terminal tree (K) and the density of CtBP2 sites per CBC6 axon terminal tree (L). There is no significant difference in the number (WT vs. rd1: p = 0.33) or density (WT vs. rd1: p = 1) of ribbon synapses made by an individual CBC cell onto an ON α-RGCs in rd1 vs. WT retinas. WT: N=8, rd1: N=8. Scale bars: A (50 µm); B-E, H-J (5 µm). All statistical comparisons were made with PWT with FDR correction.

Confocal microscope images from the IPL show CBC6 terminals overlapping with ON α-RGC dendrites (Fig. 4B). To restrict our analysis to functional synapses, we examined the overlap between ON α-RGC dendrites and ribeye-labeled puncta (CtBP2) within CBC6 terminals (Fig. 4C–E). We found that the number of putative synapses onto the ON α-RGC was the same in WT and rd1 retinas (Fig. 4F). The number of synapses per unit area of the ON α-RGC was also the same (Fig. 4G). By focusing through the depth of the IPL, we reconstructed the terminal tree of an individual CBC6 cell, which is displayed as a flattened 2D projection (Fig. 4H). Imaging ribeye puncta overlapping with the terminal tree revealed the total number of presumptive ribbon synapses formed by that CBC6 cell (Fig. 4I,J). Using this approach, we found no difference in the number of CBC6 cell output synapses between WT and rd1 retinas (Fig. 4K). Additionally, there was no difference in the density of ribbons per axon terminal (Fig. 4L). We conclude that there is no difference in the number of presumptive CBC6 synapses between WT and rd1 retina.

### Electrically evoked glutamate release from BC terminals is reduced by photoreceptor degeneration

The described results suggest that decreased synaptic efficacy from BC to postsynaptic cells in the photoreceptor-degenerated retina is caused by decreased release of the neurotransmitter glutamate. To directly confirm reduced glutamate release, we used the genetically encoded glutamate indicator iGluSnFR, delivered to RGCs with an AAV vector that transduces diverse types of RGCs indiscriminately. After allowing 2-3 weeks for viral transduction and gene expression, we isolated and mounted the retina in a chamber that enables simultaneous trans-retinal electrical stimulation and 2-photon imaging (Fig. 5A). By focusing on different depths through the inner retina, we obtained virtual cross-sections of the IPL, which showed similar baseline levels of iGluSnFR expression in WT and rd1 retinas (Fig. 5B). We placed electrodes across the thickness of the retina to enable electrical stimulation of all cells. To examine neurotransmitter release from BCs output synapse in isolation, we pharmacologically blocked all input synapses to BCs. Under these conditions, we found that a brief electrical stimulus (10 ms) evoked a transient increase in iGluSnFR fluorescence in the ON-layer of the IPL that was smaller in rd1 than in WT (Fig. 5C). The peak change in fluorescence over background (ΔF/F_0_) was 75% smaller in rd1 than in WT (Fig. 5D,E). We were concerned that retinas lacking photoreceptors might have altered tissue resistance, which could impact the effectiveness of electrical stimulation of BCs. To address this point, we varied electrical stimulus strength to generate a range of responses, from no detectable iGluSnFR signal to a saturating signal (Fig. 5F). We found that while the magnitude of the response to all stimuli (including the saturating stimulus) was lower in rd1, the sensitivity was the same, indicating that the reduced response could not be attributed to less effective electrical stimulation (Fig. 5F). Furthermore, peak iGluSnFR ΔF/F in response to application of 10 mM glutamate was no different in WT and rd1 retinas, ruling out the possibility of impaired iGluSnFR expression in rd1 retina (Fig. 5G). Taken together, these results suggest that reduced synaptic efficacy in degenerated retina is caused by reduced evoked glutamate release from ON-BCs.

**Figure 5.**
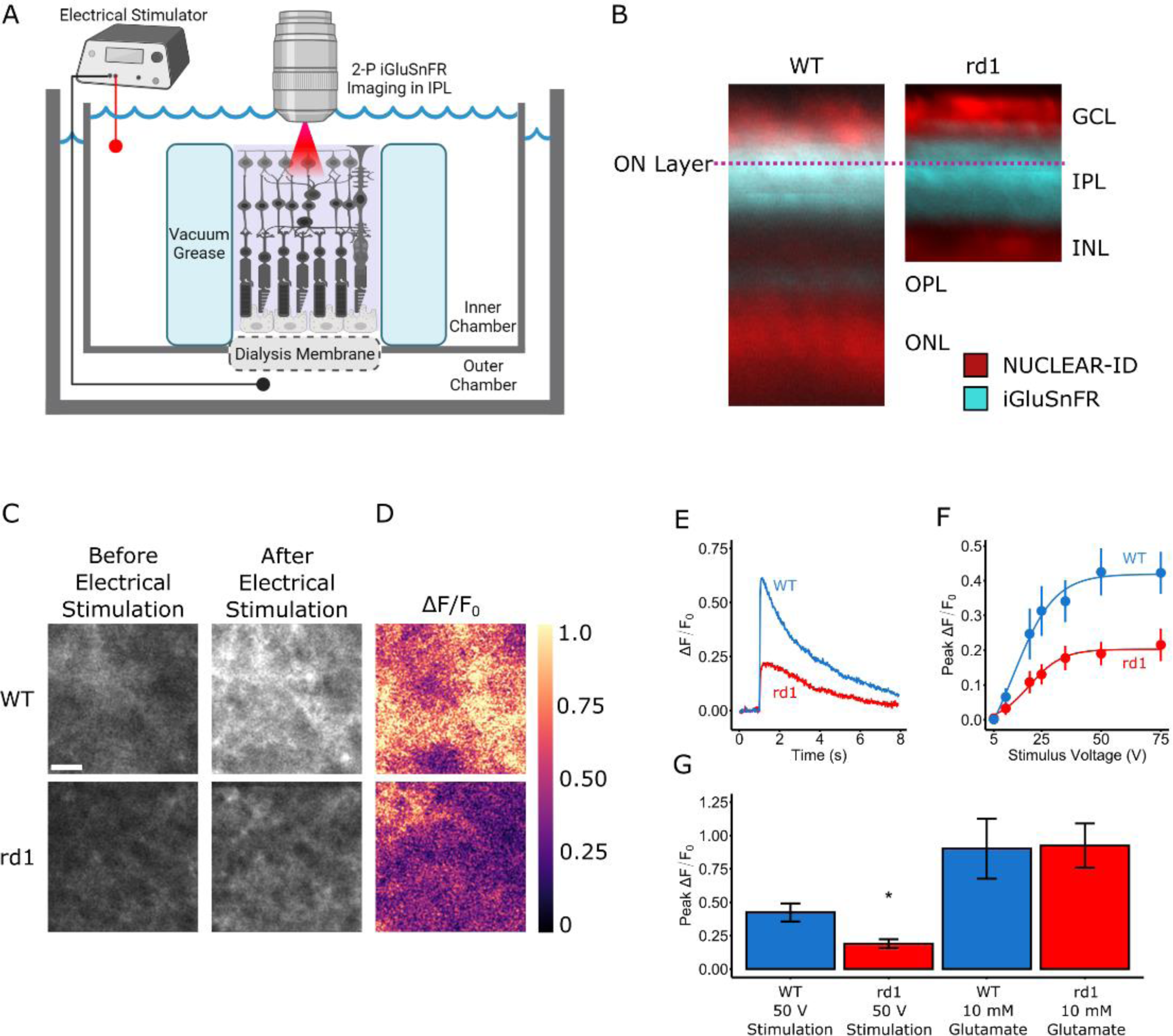
Synaptic release of glutamate from ON-BCs is reduced in retinas with photoreceptor degeneration. A) Apparatus for simultaneous trans-retinal electrical stimulation and 2-photon imaging of glutamate, the BC neurotransmitter. B) Virtual cross-sections (z-stacks) of retinas imaged with 2-photon microscopy. The genetically encoded fluorescent glutamate indicator iGluSnFR (cyan) was virally expressed throughout the IPL of WT) and rd1 retinas. The dashed line represents the ON Layer imaging plane, 10 µm into the IPL from the GCL border. Cell body layers were counterstained with a DNA-binding dye (NUCLEAR-ID, red). C) An optical section within the ON layer of the IPL before and 15 ms after stimulation, consisting of a 50 V, 2.5 ms shock in WT and rd1. D) The change in iGluSnFr fluorescence over background (ΔF/F), proportional to the change in glutamate concentration. Note that the fluorescence change upon stimulation is smaller in rd1 than in WT. E) Time course of the fluorescence change elicited by a single 2.5 ms, 50 V shock in a WT and rd1 retina. F) Peak iGluSnFR ΔF/F in response to varying shock strength. G) Peak iGluSnFR ΔF/F in response to a saturating 50 V, 2.5 ms shock is lower in rd1 than in WT (p = 0.02, N = 7 retinas). Peak iGluSnFR ΔF/F in response to application of 10 mM glutamate is no different between WT and rd1 retinas (p = 0.71, N = 7 retinas). Barplots indicate median value; error bars indicate ± SEM. Scale bar: 5 µm. All statistical comparisons were made with PWT with FDR correction.

### Retinoic acid mediates physiological remodeling of BCs

RA triggers physiological remodeling of RGCs in the degenerated retina. To explore whether RA also underlies physiological remodeling of BCs we injected the eye with BMS 493, an inhibitor of the RA receptor, RAR (Germain et al., 2009). Because the effects of RA are mediated by changes in gene expression that require days to take effect, BMS 493 treatment was given 5-7 days before functional analysis. First, we used patch clamp recording to measure voltage-gated Ca^2+^ current from BCs in retinal slices (Fig. 6A). BMS 493 increased the Ca^2+^ current by up to 3-fold in both CBC6 cells from rd1 mice and in RBCs from rd10 mice, approximating the Ca^2+^ current observed in BCs from WT retinas (Fig. 6B–E). In contrast to rd1 CBC6 cells, Ca^2+^ current in WT CBC6 cells was unaffected by BMS 493 (Fig. 6F–G). These findings suggest that elevated RA decreases the Ca^2+^ conductance and that blocking RA signaling restores the conductance, by changing the number of active Ca^2+^ channels, or altering their functional properties. Steady-state I vs. V curves show that BMS 493 caused no shift in voltage-dependent activation (Fig. 6C,E,G), suggesting that RA caused no change in gating of the Ca^2+^ channels.

**Figure 6.**
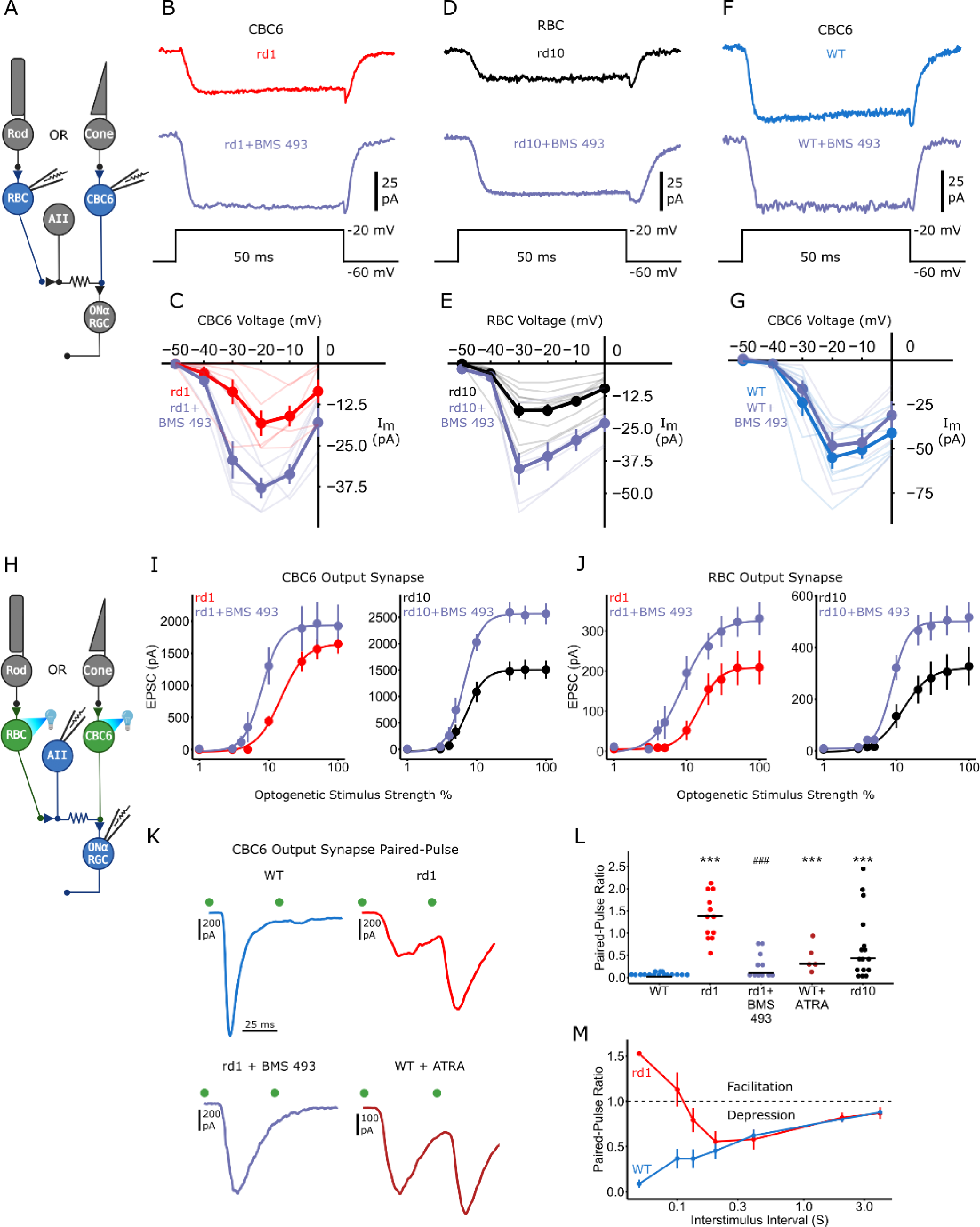
RA signaling is necessary and sufficient for physiological remodeling of BCs. A) Retinal circuit diagram showing direct recording of calcium current from either CBC6 or RBC. B,D,F) Inward calcium currents in BCs treated with BMS 493 from rd1 CBC6, rd10 RBCs, and WT CBC6 respectively. Current was activated with a depolarizing voltage step (50 ms) from −60 mV to −20 mV. Steady-state current vs. voltage (I-V) curves, elicited with a series of 10 mV incrementing depolarizing steps from −60 mV. C,E,G) I-V curves show that the BMS 493 significantly rescues voltage-gated calcium current in both CBC6 cells in rd1 retina (p <0.001) and RBCs in the rd10 retina (p <0.001). CBC6 cells: rd1, N = 6; rd1 + BMS 493, N = 5. RBCs: rd10, N = 10, rd10 + BMS 493, N = 7. BMS 493 treatment had no effect on voltage-gated calcium current in WT CBC6 cells (p = 0.93). WT: N = 8; WT + BMS 493: N = 5.) BMS 493 treatment compared to dataset from Fig.1. H) Retinal circuit diagram showing optogenetic stimulation of either CBC6 cells or RBCs. I, J) EPSC magnitude vs. optogenetic stimulus strength. BMS 493 increased EPSC magnitude in ON α-RGCs in rd1 (p=0.006) and rd10 retinas (p<0.001). RGCs: rd1, N = 15; rd1 + BMS 493, N = 11; rd10, N = 11; rd10 + BMS 493, N = 8. H). BMS 493 also increased EPSC magnitude in AII amacrine cells in rd1 (p=0.001) and rd10 retinas (p<0.001). AII amacrine cells, rd1: N=10, rd1 + BMS 493: N=9, rd10: N=14, rd10 + BMS 493: N=12. BMS 493 treatment compared to dataset from Fig.2 K) EPSCs from ON α-RGCs in response to paired pulse flashes given with a 50 ms interval. Green dots indicate light flash. WT shows complete paired-pulse depression and rd1 shows no paired-pulse depression. Note that BMS 493 induces paired-pulse depression in rd1 and ATRA eliminates paired-pulse depression in WT. L) Quantification of the paired pulse ratio (EPSC 2 / EPSC 1) for the 50 ms interstimulus interval. Compared to WT, paired EPSCs exhibited less synaptic depression in rd1 (p<0.001), rd10 (p=0.004), and ATRA-treated WT retinas (p<0.001) denoted with ***. BMS 493 treatment increased synaptic depression in rd1 retinas (p<0.001) denoted with ###. WT: N=16, rd1: N=13, rd10: N=16, rd1 + BMS 493: N=11, WT + ATRA: N=5. Horizontal lines indicate median values: WT=0.02, rd1=1.38, rd10=0.43, rd1 + BMS 493=0.10, WT + ATRA=0.30. M) Paired-pulse ratio changes as a function of interstimulus interval. Paired-pulse optogenetic pulse width: 1 ms. Error bars indicate ± SEM. Transparent traces indicate individual cells. All statistical comparisons were made with PWT with FDR correction.

Next, we examined the effect of inhibiting RA signaling on the synaptic output of BCs (Fig. 6H). BMS 493 increased optogenetically-elicited synaptic transmission from CBC6 cells by up to 35% in rd1 retina and 45% in rd10 retina (Fig. 6I), nearly completely reversing the remodeling deficit (see Fig. 2C). BMS 493 also increased synaptic transmission from RBCs, by up to 30% in rd1 retina and 50% in rd10 retina (Fig. 6J), again reducing the remodeling deficit (see Fig. 2I).

We observed that short-term synaptic plasticity is also altered in degenerated retinas. In WT retinas, the CBC6 output synapse exhibits profound paired-pulse depression, in which the first optogenetic EPSC is much larger than the second. This is consistent with WT BCs having a large Ca^2+^ current that depletes the readily-releasable pool of synaptic vesicles, leaving few available to be released by a second stimulus (Fig. 6K,L) (Gomis et al., 1999; Singer and Diamond, 2006). In contrast, paired-pulse depression is absent from rd1 or rd10 retinas, replaced in CBC6 synapses with paired-pulse facilitation, where the second response is larger than the first (Fig. 6K,L). The absence of synaptic depression can be explained by BCs in degenerated retina having a smaller Ca^2+^ current, which evokes little release during the first stimulus, preserving more vesicles to be released during a second one. The emergence of synaptic facilitation is consistent with previous results showing that reducing the size of intracellular Ca^2+^ transients in cone bipolar cells, either by reducing extracellular Ca^2+^ concentration (Borst et al., 1995) or by increasing intracellular Ca^2+^ buffer (von Gersdorff and Matthews, 1997) converts paired-pulse depression to paired-pulse facilitation.

We next asked whether the short-term plasticity profile observed in degenerated retinas could be reversed by inhibiting RA signaling. Treatment with BMS 493 converted the CBC6 output synapse in rd1 retinas from facilitating to depressing, similar to WT (Fig. 6K,L). Injecting all-trans RA (ATRA) into WT eyes reduced the amplitude of optogenetically-evoked EPSCs and reduced paired-pulse depression, suggesting that RA signaling is sufficient to trigger changes in the BC output synapse (Fig. 6K,L). These findings implicate RA as the key signal that alters short-term plasticity of the BC output synapse as a consequence of degeneration.

We further explored short-term plasticity by systematically changing the interstimulus interval. The CBC6 output synapse in WT retinas exhibited paired-pulse depression across a wide range of intervals (Fig. 6M). However, in rd1 retinas, the synapse exhibited facilitation at short interstimulus intervals, but depression at longer intervals. The involvement of multiple Ca^2+^-dependent events occurring at different subcellular locations could explain this shift in short term plasticity. With a smaller Ca^2+^ current, intracellular Ca^2+^ will rise more slowly and be more tightly constrained within a nanodomain near the plasma membrane (Zucker and Regehr, 2002), preferentially engaging different Ca^2+^ sensors than a larger Ca^2+^ current. Different isoforms of synaptotagmin, each with a distinct Ca^2+^ affinity, underlie different events in synaptic release (e.g. vesicle replenishment and fusion) (Bacaj et al., 2013; Jackman et al., 2016; Weingarten et al., 2022) and these Ca^2+^ sensors will be differentially activated depending on the intracellular Ca^2+^ profile, which is sensitive to the interval between individual Ca^2+^ transients.

### Physiological remodeling of BCs alters spatiotemporal information processing

In the WT retina, sensory information from an array of photoreceptors signals through a series of retinal neurons, ultimately converging onto an RGC to produce a classical receptive field. In a photoreceptor-degenerated retina with the responses of a particular type of BC under optogenetic control, each RGC receives a projective field, reflecting the convergence of information from that specific BC cell type. Here we compare rd1 and WT retinas to ask whether photoreceptor degeneration alters the CBC6 to ON α-RGC projective field. We optogenetically stimulated small clusters of CBC6 cells with 50 µm spots of light to yield a spatial map of BCs projecting onto an individual ON α-RGC (Fig. 7A–C). We found that the spatial map in rd1 retinas is nearly identical to the map in WT retina, but scaled down linearly (Fig. 7D,E), presumably owing to decreased synaptic transmission. This finding is consistent with previous work showing little change in cone-mediated RGC receptive field size following rod degeneration, in a different mouse model of RP (Scalabrino et al., 2022).

**Figure 7.**
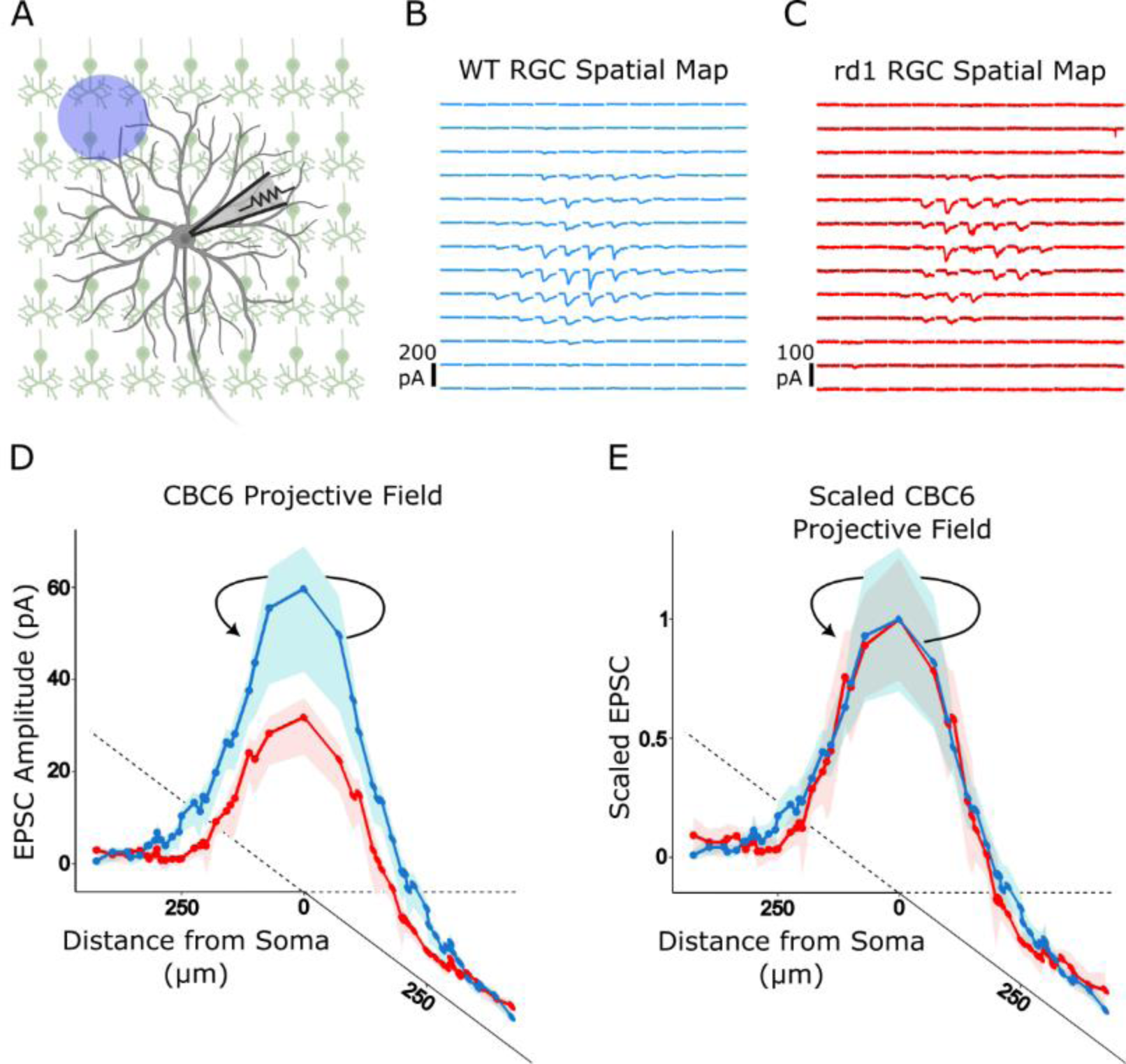
The spatial map of the BC to RGC network is preserved in photoreceptor degenerated retina but scaled down linearly. A) Schematic diagram of the experiment. To map the projective field corresponding to many CBC6 cells converging onto a single ON α-RGC, we used 50 µm non-overlapping spots of light to optogenetically stimulate clusters of BCs, covering the entire dendritic tree of the ON α-RGCs. B-C) Example spatial projective field map from a WT (B) and rd1 (C) retina. Detectable EPSCs were elicited over ∼200 µm^2^ of the 650 µm^2^ mapped area. D) Group data from ON α-RGCs (N=9 each), with the projective field represented as mean EPSC vs. distance from the cell soma. Data from all directions from the cell center were averaged and are shown duplicated for clarity. Note that EPSCs were smaller in rd1 than WT (p=0.012), but there was no difference in spatial weighting of the linearly scaled projection field (p=0.47). Variability ribbons indicate 95% confidence intervals. All statistical comparisons were made with PWT with FDR correction.

We next compared the temporal properties of the CBC6 to ON α-RGC network in WT and photoreceptor degenerated retinas. We optogenetically stimulated populations of CBC6 cells at different rates and recorded either EPSCs in whole-cell patch recordings or spiking in cell-attached patch recordings from individual RGCs (Fig. 8A). In the WT retina, a low frequency train of stimuli (0.25 Hz) evoked a non-decrementing train of EPSCs (Fig. 8B,G). Even with higher frequency stimulation (up to 7.5 Hz), EPSCs followed reliably, although their amplitudes declined during the train (Fig. 8B,G–I). In rd1 and rd10 retinas, EPSCs were non-decrementing at low frequency (0.25 Hz), but their amplitudes declined rapidly at higher frequencies, with EPSCs failing completely by the end of the 7.5 Hz stimulation train. To confirm that optogenetic stimulation caused reliable depolarizations that were equivalent in WT and rd10 retinas, we recorded from CBC6 cells under current clamp while applying trains of light flashes. Except for a ∼50% decline in amplitude between the first and second depolarization, subsequent depolarizations showed little decrement for at least 3 s, when flashes were repeated at 7.5 Hz (Fig. 8C,D). Hence changes in presynaptic depolarizations cannot account for the failure of postsynaptic responses.

**Figure 8.**
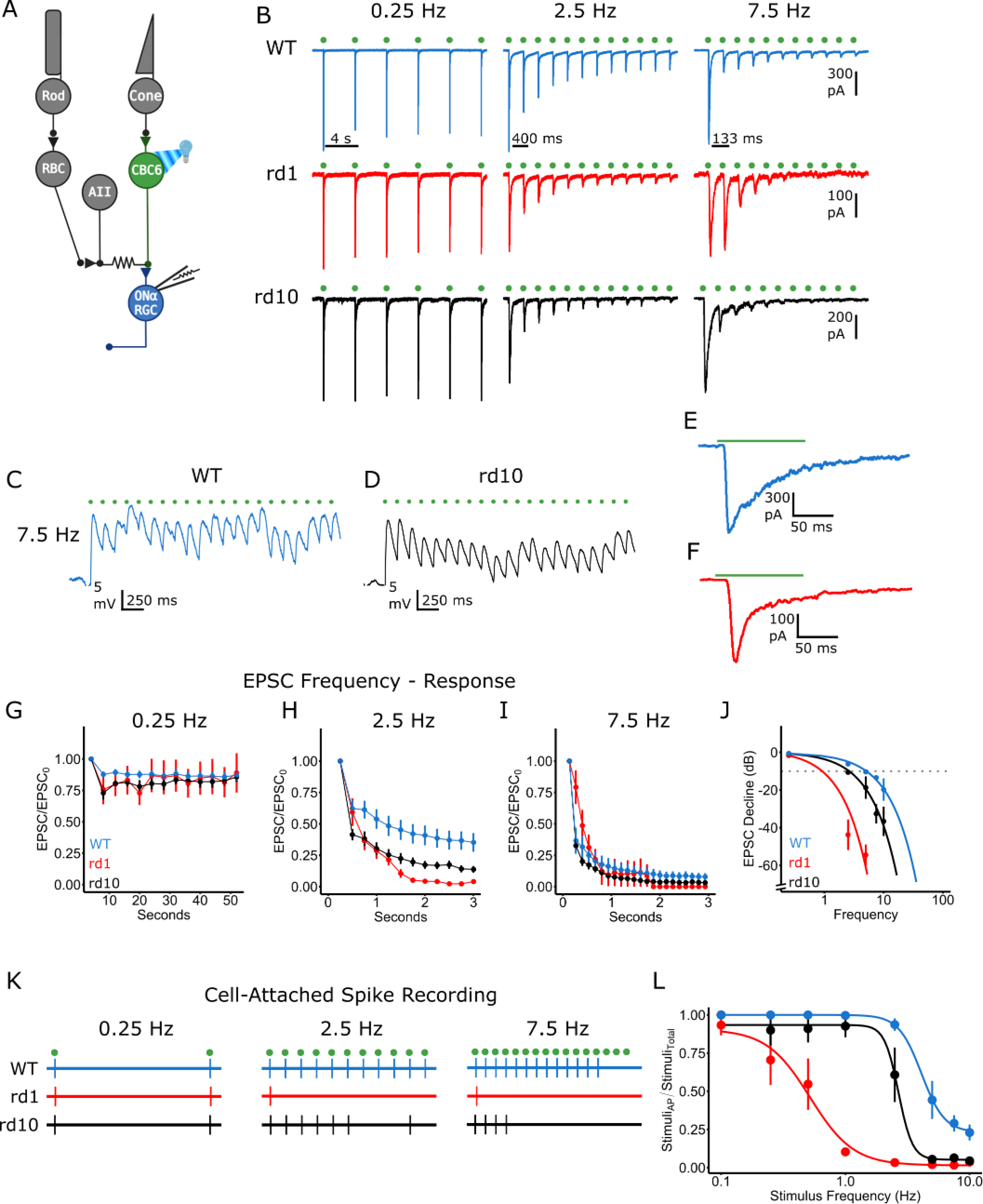
Photoreceptor degeneration reduces the frequency response in the cone pathway. A) CBC6 cells were optogenetically stimulated at different rates while EPSCs were recorded in an ON α-RGC. B) EPSCs elicited by stimuli repeated at 0.25 Hz, 2.5 Hz, and 7.5 Hz. Green dots indicate the time of light flash: 2.5 ms pulse width. C, D) Recording membrane voltage directly in CBC6 from WT and rd10 little decrement in depolarization following 7.5 Hz optogenetic stimulation. E, F) Immediately following a 5 Hz optogenetic stimulation train, a 100 ms pulse was given resulting in a large EPSC despite a depressed synapse in both a WT and rd1. G – I) EPSCs in rd1 and rd10 retinas diminish faster with increasing stimulation frequency. J) EPSC decline expressed in dB 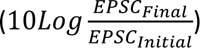 for each stimulation frequency. The continuous line is the fit with a linear function. (WT: N=7, *y* = −0.143 − 1.93*x*, R^2^= 1.00; rd1: N = 9, *y* = 1.42 − 12.5*x*, R^2^= 0.99,; rd10: N=10 *y* = 0.272 − 3.93*x*, R^2^= 0.95). The dotted line is *y* = −10 *dB*. EPSCs for rd1 at 7.5 and 10 Hz become immeasurably small at steady state. K) Raster of cell-attached spike recordings from stimulation at 0.25 Hz, 2.5 Hz, and 7.5 Hz. Green dots indicate light flash. L) The ratio of stimuli that produced an action potential to the total number of stimuli per frequency. Compared to WT, rd1 and rd10 ON α-RGCs successfully encoded fewer stimuli as stimulation frequency increased. WT: N=6, rd1, N=6, rd10, N = 5 cells.(rd1: p<0.001, rd10: p=0.034. PWT with FDR correction). Error bars indicate ± SEM.

Comparing across all stimulation rates, the frequency at which EPSCs declined by 90% (−10 dB) at steady state was 5.25 Hz for WT, 0.68 Hz for rd1, and 2.47 Hz for rd10. (Fig. 8J). Despite this dramatic decline, an EPSC could still be evoked if a greatly prolonged depolarizing stimulus were applied at the end of the train (Fig. 8E,F), suggesting that even though the pool of readily-releasable vesicles had been depleted, it could be resupplied from a reserve pool, given a sufficiently large and prolonged intracellular Ca^2+^ signal. Vesicle resupply is a Ca^2+^-accelerated processes, involving an isoform of synaptogamin distinct from the isoform that underlies vesicle fusion (Weingarten et al., 2022) and perhaps the large Ca^2+^ signal engages this process.

Decrementing synaptic efficacy would be expected to adversely impact action potential reliability in RGCs. To test this, we made cell-attached patch-clamp recordings to non-invasively monitor spike activity from ON α-RGCs. In the WT retina, RGC firing followed optogenetic BC stimulation reliably up to 7.5 Hz (Fig. 8K), consistent with temporal tuning of normal light responses in ON α-RGCs in WT retinas (Tien et al., 2017). In the rd1 and rd10 retina, firing failed to follow even at 2.5 Hz stimulation frequency. We compiled firing reliability data over a wide range of stimulus frequencies and train durations. RGCs failed to follow BC stimulation at a low frequency in rd10 and an even lower frequency in rd1 (Fig. 8L). The decline of reliability in the CBC6 to ON α-RGC network can be attributed entirely to changes in the BCs. We found no difference in the frequency response of ON α-RGC to firing induced by direct current injection in WT and degenerated retinas (data not shown). These findings illustrate the severe deficit that BC physiological remodeling imposes on information transfer to RGCs in the degenerated retina.

Equivalent experiments on RBC output synapse showed an even more extreme frequency-dependent decline in photoreceptor degenerated retina (Fig. 9A,B). Even at 0.25 Hz, EPSCs from rd1 and rd10 retinas began decrementing while WT did not (Fig. 9B,C). Higher stimulation frequency led to the near complete loss of EPSCS in rd1 and rd10 at steady state (Fig. 9B,E–G). Current clamp recordings from RBCs confirmed that optogenetic stimulation caused reliable depolarizations that were equivalent in WT and rd10 retinas (Fig. 9C,D). Hence as was the case for CBC6 cells, changes in presynaptic depolarizations in RBCs cannot account for the failure of postsynaptic responses.

**Figure 9.**
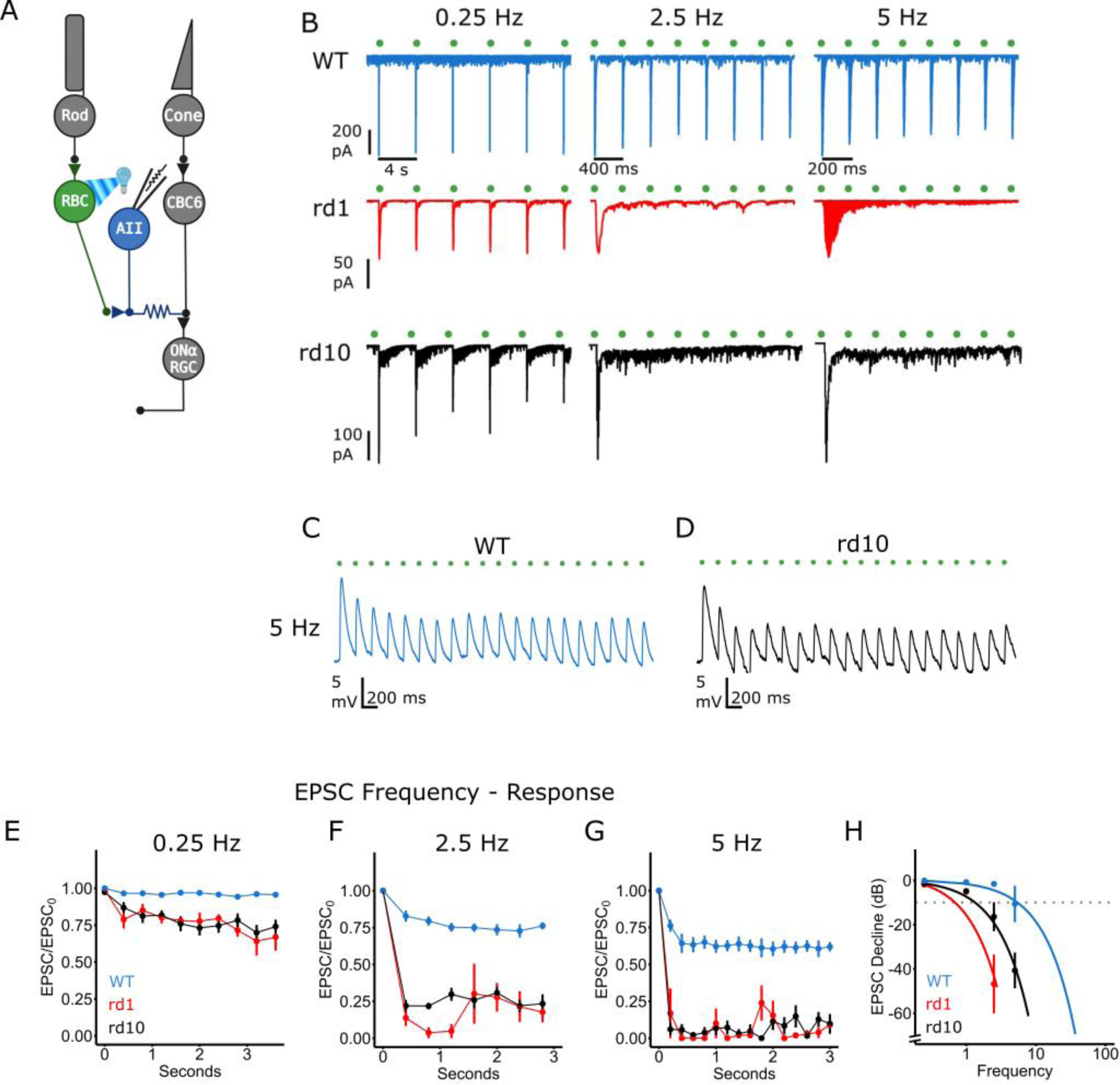
Photoreceptor degeneration reduces the frequency response in the rod pathway. A) RBCs cells were optogenetically stimulated at different rates while EPSCs were recorded in an AII amacrine cell. B) EPSCs elicited by stimuli repeated at 0.25 Hz, 2.5 Hz, and 5 Hz. Green dots indicate the time of light flash: 5 ms pulse width. C, D) Recording membrane voltage directly in RBCs from WT and rd10 little decrement in depolarization following 5 Hz optogenetic stimulation. E–G) EPSCs in rd1 and rd10 retinas diminish faster with increasing stimulation frequency. H) EPSC decline expressed in dB 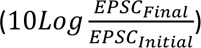 for each stimulation frequency. The continuous line is the fit with a linear function. (WT: N=6, *y* = −0.28 − 1.89*x*, R^2^= 0.90; rd1: N=5, *y* = 2.63 − 18.4*x*, R^2^= 0.99,; rd10: N=11, *y* = 0.579 − 7.91*x*, R^2^= 0.94). The dotted line is *y* = −10 *dB*. EPSCs for rd1 at 5 Hz become immeasurably small at steady state. Error bars indicate ± SEM.

Comparing all stimulation rates, the frequency at which EPSCs declined by 90% (−10 dB) at steady state was 5.14 Hz for WT, 0.69 Hz for rd1, and 1.34 Hz for rd10. (Fig. 9H). Since the vast majority of ON BC types receive direct or indirect input from RBCs, the decrementing responses in photoreceptor degenerated retinas is expected to adversely impact a wide variety of ON-RGCs beyond just the ON α-RGC subtype.

## Discussion

This paper reports the surprising finding that photoreceptor degeneration leads to dramatic functional weakening of the output synapses of ON-BCs. RBCs, the most numerous type of BC in the rod-dominant mouse retina, and CBC6 cells, which receive information from both rod and cone pathways, are both affected. The net effect of these changes in BCs is to reduce the amplitude and narrow the frequency response of light onset signals sent to RGCs and onward to the brain. Our imaging results with iGluSnFr suggests that glutamate release from OFF-BCs might also be reduced in the degenerated retina, which might also reduce signals at light offset. New cell-type selective promoters for particular OFF-BCs could help address this point.

Previous experiments on a rat model of RP showed that ON-RGCs exhibit reduced spontaneous firing, paralleling the loss of the RGC light response (Sekirnjak et al., 2011). The most obvious explanation for the decline in spontaneous and evoked ON-RGC activity is that the diminishing number of photoreceptors drives a declining amount of excitatory synaptic input. However, the possibility that changes intrinsic to ON-BCs also contributed to changes in ON-RGC activity remained unexplored. Our discovery that the output synapse of ON-BCs is dramatically weakened adds an important new mechanism that may contribute to changes in ON-RGC activity across species, not only corrupting retinal signal processing, but also contributing to the pathophysiology of vision loss.

Another consequence of photoreceptor loss is the emergence of membrane potential oscillations in the inner retina, ultimately resulting in spontaneous burst firing in RGCs (Trenholm and Awatramani, 2015). The oscillation is initiated in the electrically coupled network of AII amacrine cells (Choi et al., 2014), but it remains to be determined if RA-mediated events are involved.

### Mechanism of physiological remodeling

Our findings suggest that RA is necessary and sufficient for physiological remodeling of the ON-BC output synapse. Adding exogenous ATRA mimics changes in synaptic release that are characteristic of degenerated retinas (Fig. 6K). Inhibiting RAR with BMS 493 reverses changes in synaptic release and restores voltage-gated Ca^2+^ current to near WT levels (Fig. 6). Our previous studies utilizing an RA-induced reporter demonstrated elevated RA-induced gene expression in degenerated retina (Telias et al., 2019).

It remains unclear where the excessive RA is produced. RA has been implicated in physiological remodeling not only in the output synapses of BCs but also in RGCs (Telias et al., 2019). Moreover, RA is critical for morphological remodeling of BC dendrites, at their input synapses (Lin et al., 2012). The fact that multiple cell types are affected in a degenerated retina suggests that RA might act in a paracrine fashion, produced by one cell type (perhaps RPE cells), and diffusing throughout the retina to trigger remodeling in many cell types. Immunocytochemistry and enzyme assays show that the retina, choroid, and RPE all express isoforms of RALDH, the enzyme that synthesizes RA (Harper et al., 2015). In the RPE, RALDH normally converts all-trans retinaldehyde released from photobleached rhodopsin into RA (McCaffery et al., 1996). The loss of photoreceptors might increase the concentration of retinaldehyde, leading to increased production of RA by the RPE. In this scenario, RA would be an intercellular signal, diffusing from the outer to the inner retina to exert its actions. Alternatively, it is possible that RA is synthesized within particular retinal neurons and serves as an intracellular messenger. In neurons in the hippocampus, local production of RA in dendrites regulates local gene translation, a key event underlying homeostatic synaptic plasticity (Thapliyal et al., 2022).

How does elevated RA decrease voltage-gated Ca^2+^ current and synaptic transmission? The size of the Ca^2+^ current in ON-BCs is reduced by photoreceptor degeneration, but the activation and inactivation properties are unchanged, suggesting a decreased level of channel protein rather than a change in channel type. RAR is an enhancer of gene transcription, which often results in increased expression of specific proteins. However, there are many examples of RA-induced down-regulation of proteins, mediated by a sign-inverting gene regulatory cascade (Balmer and Blomhoff, 2002).

L-type Ca^2+^ channels are complexed with accessory proteins, including alpha-2/delta subunits that determine Ca^2+^ current density and activation/inactivation kinetics (Dolphin, 2009), in addition to controlling vesicle release probability at synapses (Hoppa et al., 2012). It is possible that functional down-regulation of Ca^2+^ channels is caused by misregulation of alpha-2/delta, as occurs in sensory neurons in dorsal root ganglia after nerve injury (D’Arco et al., 2015). Transcriptomic analysis of BCs indicates that there are changes in gene expression, but they are limited to only a few genes(Gilhooley et al., 2021), which could simplify the task of identifying the molecular basis of calcium current down-regulation. Many other RA-induced biochemical events could also result in decreased Ca^2+^ current, including altered post-translational modification or altered membrane trafficking of Ca^2+^ channel proteins (D’Arco et al., 2015).

Is the loss of Ca^2+^ channels sufficient to account for the decline in synaptic release, without invoking additional changes in the release machinery? We find no decline in the number of synaptic contacts between BCs and RGCs and no loss of synaptic ribbons, a key component of the BC synaptic machinery. Hence, at least at this age (p60) in rd1 mice, photoreceptor degeneration causes no obvious structural changes in the BC output synapse. We did observe one interesting phenomenon concerning the frequency-dependence of the ON-BC output synapse in the degenerated retina, but it also may be explained by decreased Ca^2+^ influx during stimulation. We found that paired-pulse depression was dramatically decreased in rd1 and rd10 retinas (Fig. 6K), whereas there was a great increase in failure of the synapse with more sustained, high-frequency stimulation (Fig. 8J). Intracellular Ca^2+^ regulates synaptic vesicle fusion to the plasma membrane, but it is also thought to regulate other steps in the synaptic vesicle cycle, such as vesicle docking and priming (Wan and Heidelberger, 2011). In fact, there is evidence for multiple Ca^2+^-dependent steps in the vesicle cycle at BC ribbon synapses (Gomis et al., 1999). If Ca^2+^ influx were sufficient to cause vesicle fusion but insufficient to accelerate vesicle resupply, repetitive stimulation might deplete the pool of readily releasable vesicles before they could be replenished. Different isoforms of synaptotagmin mediate fusion and resupply (Weingarten et al., 2022), providing opportunities for understanding how reduced Ca^2+^ influx in remodeled BCs affects synaptic transmission at the molecular level.

### Functional impact

Our results imply that physiological remodeling of ON-BCs will constrain information processing further along in the visual system, limiting visual perception. The deficit in temporal coding caused by decreased ON-BC output was particularly severe, which is expected to have a particularly large impact on perception of object motion. Humans with RP often fail to detect moving objects or report motion in the opposite direction of actual movement (Turano and Wang, 1992), even if they retain significant visual acuity for detecting static objects (Alexander et al., 1998). RP is a progressive disease that causes gradual vision loss, but the distinct contributions of decreased photoreceptor density vs. physiological remodeling to vision decline is unclear. Our previous results (Telias et al., 2022) showed that treatments that reverse physiological remodeling could substantially rescue visual contrast sensitivity in rd10 mice at 2-3 months of age, when photoreceptor degeneration is already severe. We now know that physiological remodeling applies not only to RGCs, but also to BCs, but whether remodeling of one cell type or the other predominates in altering visual perception is also unknown.

In certain respects, the physiological remodeling that we characterized in ON-BCs might be considered a homeostatic mechanism, analogous to other compensatory events caused by partial photoreceptor loss (Lee et al., 2022). In the healthy retina, photoreceptors tonically release glutamate, which keeps ON-BCs hyperpolarized. Hypothetically, by eliminating glutamate release, photoreceptor death should depolarize the ON-BCs, chronically activating their voltage-gated Ca^2+^ channels, and increasing their tonic release of glutamate onto RGCs. If this chronic synaptic excitation of RGCs were sufficient to trigger firing, it could impair encoding of synaptic information from other BCs that had yet to undergo remodeling. One way that ON-BCs could compensate for excessive depolarization is by down-regulating their Ca^2+^ channels. While the loss of Ca^2+^ channels might be adaptive, by preventing excessive output from ON-BCs, it comes at a cost. We have demonstrated that physiological remodeling limits the dynamic range of the synaptic output of CBC6 cells (Fig. 2C) and RBCs (Fig. 2I) and severely limits temporal reliability (Fig. 7,8), which is maladaptive for encoding visual information. The degree of photoreceptor loss may determine whether physiological remodeling is adaptive or maladaptive.

Our findings that physiological remodeling of ON-BCs, as well as RGCs, can be reversed by inhibiting RA has important implications for preserving sight in low-vision patients with RP and perhaps other photoreceptor degenerative disorders. They also have important implications for vision restoration in advanced RP patients with nearly complete photoreceptor loss and little remaining light perception. One treatment approach for these patients is to install light responses in surviving neurons of the outer retina, including ON-BCs, either through optogenetic stimulation (Lagali et al., 2008; Sahel et al., 2021) or electrical stimulation with sub-retinal implants (Roux et al., 2016). While ON-BCs in mouse models of RP do retain the ability to transmit optogenetically-elicited synaptic signals to RGCs (Lagali et al., 2008), our findings suggest that physiological remodeling of the ON-BC output synapse will limit downstream information processing and visual perception. Our results indicate that reduced frequency-response of the ON-BC output synapse, seen as a failure of EPSCs, severely impairs RGC spike-rate coding. In the clinical context, this may severely limit the effectiveness of optogenetic therapies targeted to ON-BCs or upstream remnant photoreceptors in degenerated retina. The same problems may also apply to signals initiated by stem-cell derived regenerated photoreceptors inserted into the outer retina. Inhibiting RA synthesis or signaling may remove these problems and augment the effectiveness of any vision restoration technology that delivers light-elicited signals to the blind retina. RA inhibition can be accomplished with pharmacological inhibitors of RALDH or RAR (Telias et al., 2022), or with targeted AAV vectors that deliver genetic-based inhibitors of RALDH and RAR to specific retinal cell types, for example ON-BCs or RGCs (Bi et al., 2006; Lagali et al., 2008).

